# Regulated proteolysis of green fluorescent protein fusion proteins in *Aspergillus fumigatus*

**DOI:** 10.1101/2025.07.14.664774

**Authors:** Sanjoy Paul, W. Scott Moye-Rowley

## Abstract

Aspergillosis caused by *Aspergillus fumigatus* is a clinical issue of such severity that the World Health Organization has designated this organism as 1 of the 4 most critical fungi to study. Progress in *A. fumigatus* has been limited by the availability of genetic tools with which to study this filamentous fungus. Currently available means of altering the dosage of genes and gene products include construction of disruption mutants as well as regulated promoters. These are powerful techniques but somewhat limited for the analysis of essential genes. Here we describe a new method that permits regulated proteolysis of any *A. fumigatus* protein that can be made as a fusion protein to the well-described Green Fluorescent Protein (GFP) of *Aequorea victoria*. A GFP fusion protein of interest can be targeted for degradation using a single-chain antibody called a nanobody that recognizes GFP (GFPNb). This GFPNb is in turn fused to an E3 ligase protein called Rnf4 from rat that efficiently ubiquitinates target proteins. A fusion gene was constructed under control of a doxycycline-inducible promoter that produced a GFPNb-Rnf4 fusion protein in *A. fumigatus*. Here we show that production of this GFPNb-Rnf4 protein led to the rapid proteolysis of a variety of GFP fusion proteins. Additionally, we found that some GFP fusion proteins triggered a corresponding genomic response when their degradation was induced while others were simply degraded. These studies provide a new means to directly regulate protein levels in *A. fumigatus* and generate new alleles of genes exposing the underlying regulatory circuitry.

**Importance:** *Aspergillus fumigatus* is the major filamentous fungal pathogen of such importance in infectious disease that it has been designated as one of the 4 most critical fungal organisms to study. One of the limiting capabilities in the analysis of *A. fumigatus* is the limiting genetic toolbox that can be employed in this fungus. Here we describe the first system that allows the regulated degradation of a protein of interest using a recombinant single protein chain antibody (nanobody) directed against Green Fluorescent Protein (GFP). GFP is routinely used to construct fusion proteins that in turn allow localization of the fluorescent protein and immunological detection. Here we use this nanobody to deliver a ubiquitin ligase that in turn causes rapid depletion of GFP fusion proteins. This allows direct perturbation of protein levels in *A. fumigatus*, a feature not previously accessible to experimentation.

## Introduction

*Aspergillus fumigatus* is the major human filamentous fungal pathogen with a mortality approaching 80% in particular patient populations (1). Azole antifungal drugs are the clinical gold standard for treatment of this pathogen but, the appearance of resistant variants (2), coupled with their potentially high mortality puts the continued efficacy of this treatment at risk (3). These factors have contributed to aspergillosis caused by *A. fumigatus* to be listed as a fungal infection of critical importance in the recent categorization by the World Health Organization (4). Although the severity of disease associated with *A. fumigatus* is clearly appreciated, our understanding of the basic biology of this fungus is still at an early stage.

Study of *A. fumigatus* is limited by the relative lack of availability of important experimental tools. For example, promoters providing regulated expression of genes of interest have been produced and effectively permit placing gene products under control of doxycycline-inducible or -repressible transcriptional control (5, 6). Additionally, strong constitutive promoters like *gpdA* or *hspA* have been described that often lead to elevated expression of desired gene products (7–9). While these transcriptional control regions have been effectively utilized for altering gene expression, an important limitation is that this strategy acts only to control the level of mRNA of a target gene. This restricts their utility when stable proteins are being investigated. A system that directly impacted protein levels would be more likely to lead to rapid depletion of a gene product under study.

At least two different systems have been developed initially in *Saccharomyces cerevisiae* that have been seen to effectively cause degradation of a desired protein product. The first of these leveraged the receptor, called Tir1, for the plant hormone auxin (10, 11). Tir1 is able to bind to endogenous yeast components to form an SCF E3 ubiquitin ligase. Upon auxin binding, Tir1 is triggered to bind a short protein region called the auxin-inducible degron (AID) that can be fused to target proteins of interest (12). In *S. cerevisiae* (12) and two different *Candida* species (13, 14), this system works well to target gene products fused to AID for rapid protein degradation.

A second, more recently described targeted protein degradation system takes advantage of a single chain antibody provided by camelid species (15) that has been engineered to recognize the well-known green fluorescent protein (GFP) moiety that is commonly used to facilitate localization of proteins as GFP fusion proteins (16). Expression of this single chain antibody or nanobody permitting GFP recognition as a fusion with an E3 ligase component leads to ubiquitination and proteasomal degradation of a GFP fusion protein of interest (17).

We attempted to deploy the auxin-inducible degradation system in *A. fumigatus* but were faced with technical problems including the previously described growth inhibitory effects of auxin (18). These issues drove us to adopt the nanobody-based degradation of GFP fusion proteins which we found to be effective in targeting these recombinant proteins in *A. fumigatus* for rapid proteolysis. We fused the anti-GFP nanobody or GFPNb to the ubiquitin ligase enzyme provided by the human Rnf4 gene product and placed this chimeric protein under doxycycline-inducible (dox-on) promoter control (6). The addition of doxycycline triggered rapid and efficient expression of the GFPNb-Rnf4 fusion protein and attendant degradation of multiple different GFP fusion proteins located in different subcellular compartments. We believe this nanobody system provides a useful, new means to directly control levels of proteins under study in this filamentous fungal pathogen. We also contrast use of this regulated system of proteolysis to analyze the biological effects caused by depletion of the essential ergosterol biosynthetic enzyme Erg6 using doxycycline-repressible promoter.

## Results

### Strategy to express a GFP nanobody coupled to an E3 ligase enzyme in *A. fumigatus*

Our original attempts to produce a regulated proteolysis system in *A. fumigatus* focused on using the auxin-inducible degradation system that has been used in *Candida* species (13, 14). There were a number of technical issues with this including the growth inhibitory effects caused by the presence of auxin (18) and expression of the plant Tir1 E3 ligase (data not shown). We abandoned these efforts and switched to production of an anti-GFP camelid single chain antibody or GFP nanobody (GFPNb) fused to the rat E3 ligase Rnf4 (19). This construct has been previously used to target GFP fusion proteins for proteasomal degradation in mammalian cells (17). The GFPNb binds to GFP and will deliver the covalently-fused E3 ligase activity of Rnf4 to this position. We engineered the expression of this to be under the control of a doxycycline-inducible promoter in order that levels of the GFPNb-Rnf4 be low unless this compound was added to the medium (Figure 1A). We generated versions of the GFPNb-Rnf4 that were expressed as fusion proteins with the estrogen-binding domain (EBD) from the mammalian estrogen receptor previously modified to be tightly regulated in *Saccharomyces cerevisiae* (20). We previously found that this EBD domain helped repress the activity of the cre recombinase expressed in *Candida glabrata* cells (21).

**Figure 1.**
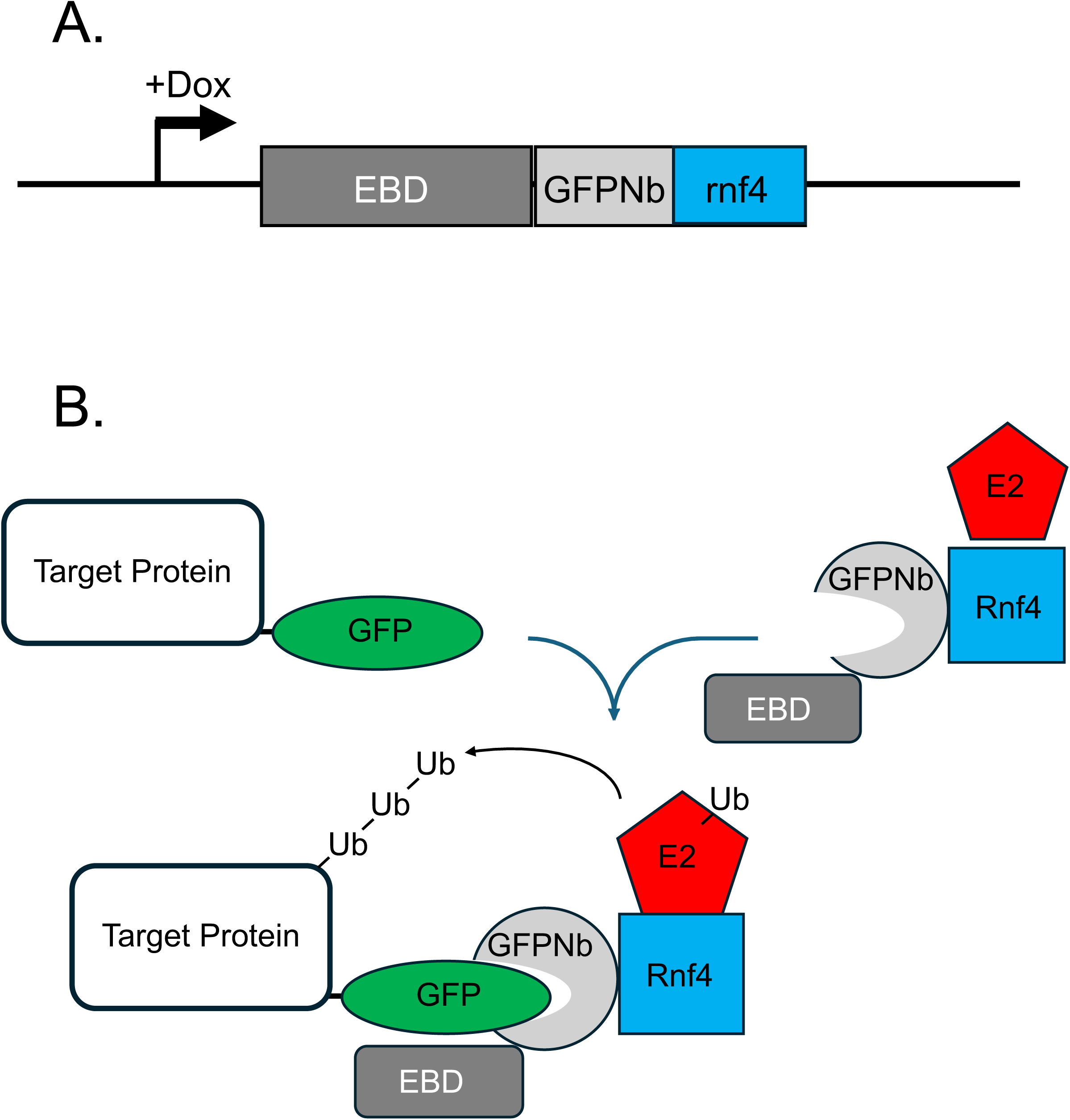
Design of nanobody-directed proteolysis system. A. Inducible expression of the anti-GFP nanobody (GFPNb) fusion protein. A doxycycline-inducible promoter was used to express the estrogen-binding domain (EBD) fused to the camelid anti-GFP nanobody and the E3 ligase fragment from the Rnf4 ubiquitin ligase. We also produced this fusion protein lacking the EBD moiety. All GFPNb-expressing clones contained a 4X HA tag at the GFP amino-terminus to allow detection of fusion protein expression. B. The theoretical function of the two different components of the GFPNb-directed proteolysis system is depicted. First, a translational fusion between a target protein of interest and GFP is generated in the *A. fumigatus* genome. This target protein-GFP fusion is susceptible to the activity of the GFPNb-containing fusion protein. The GFP segment of the target protein is recognized by the anti-GFP nanobody with its associated EBD and E3 ligase enzyme Rnf4. Rnf4 in turn recruits an E2 enzyme, ultimately leading to the formation of polyubiquitin chains on the target protein with attendant degradation by the proteasome.

The manner in which the EBD-GFPNb-Rnf4 fusion protein is believed to act is shown in Figure 1B. Substrate proteins are expressed as GFP fusions in the presence of the doxycycline-inducible EBD-GFPNb-Rnf4. Upon doxycycline addition, the nanobody-containing protein is expressed. The GFPNb delivers Rnf4 to the target GFP fusion protein. Rnf4 is able to bind to endogenous E2 proteins that in turn permit ubiquitin to be transferred by Rnf4 activity to GFP substrate proteins which are then degraded by the proteasome.

### Degradation of GFP by the EBD-GFPNb-Rnf4 chimeric protein

To test the in vivo efficacy of the EBD-GFPNb-Rnf4 fusion protein, we constructed a strain that produced an epitope-tagged form of GFP from the constitutive *gpdA* promoter. Appropriate transformants were grown in the presence (+) and absence (-) of doxycycline and β-estradiol (to activate proteins fused to the EBD (20)) and levels of EBD-GFPNb-Rnf4 and GFP assessed by western blotting using antibodies recognizing the HA or FLAG epitopes respectively (Figure 2).

**Figure 2.**
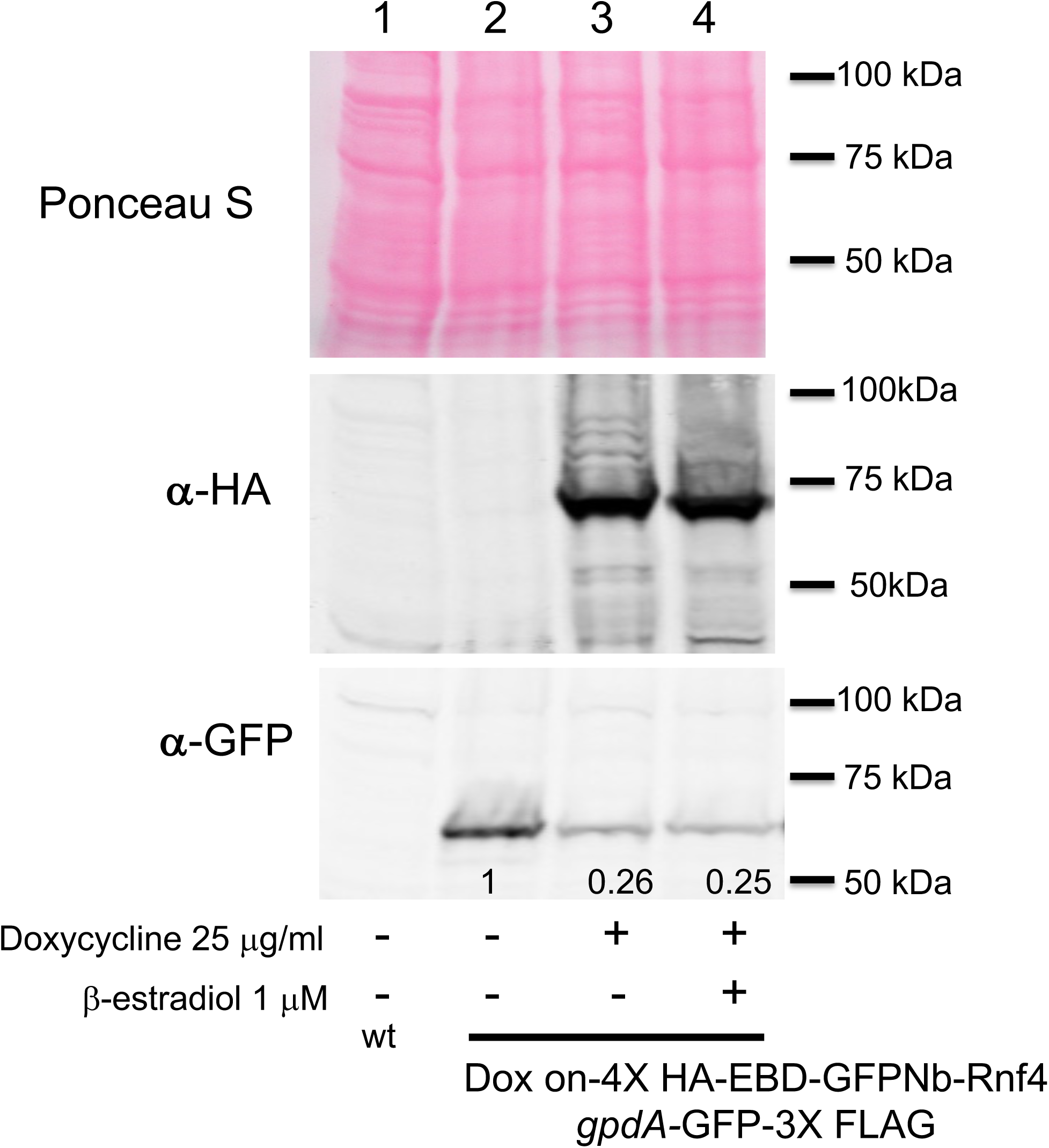
Sensitivity of *gpdA*-driven GFP to GFPNb-directed proteolysis. Whole cell protein extracts were prepared from AfS35 cells (wt) or from AfS35 containing an integrated doxycycline-inducible EBD-GFPNb-Rnf4-expressing clone and a gpdA-controlled GFP gene. Strains were grown in Sabouraud dextrose media in the absence (-) or presence (+) of doxycycline and β-estradiol as indicated. Equal amounts of protein from each extract were resolved by SDS-PAGE and analyzed by western blotting. The transferred membrane was stained with Ponceau S to confirm equal loading followed by visualization with antibodies recognizing the hemagglutinin tag (α-HA) or GFP (α-GFP). Molecular mass standards are indicated on the righthand side of each panel. The numbers at the bottom of lanes 2-4 provide quantitation of the GFP signal relative to that seen in the absence of either doxycycline or β-estradiol.

Induction of EBD-GFPNb-Rnf4 expression in the presence of doxycycline triggered a depletion of the *gpdA*-driven GFP-Flag protein. The presence of β-estradiol had no effect on the doxycycline-stimulated or unstimulated degradation of GFP-Flag suggesting that, at least in this context, the EBD was not impacted by ligand. These data support the view that the EBD-GFPNb-Rnf4 is active when expressed in *A. fumigatus* and capable of triggering the proteolysis of a cytoplasmic GFP. To begin to evaluate the range of substrates that EBD-GFPNb-Rnf4 was capable of degrading, we used a nuclear GFP fusion protein encoded by *ncaA*-GFP

### Degradation of a nuclear GFP-marked protein, NcaA-GFP

We have previously analyzed the subcellular distribution of a nuclear protein called NcaA that is a coactivator of the transcription factor AtrR (22, 23). We constructed a C-terminal fusion between NcaA and GFP that demonstrated this protein was almost exclusively localized to the nucleus (24). We introduced our doxycycline-inducible EBD-GFPNb-Rnf4 construct into this strain to allow us to assess if the nanobody-mediated proteolysis could be seen on a nuclear substrate. A strain containing the *ncaA*-GFP fusion gene with or without the EBD-GFPNb-Rnf4-expressing construct were grown in the presence or absence of doxycycline and analyzed for levels of the NcaA-GFP fusion protein using western blotting (Figure 3).

**Figure 3.**
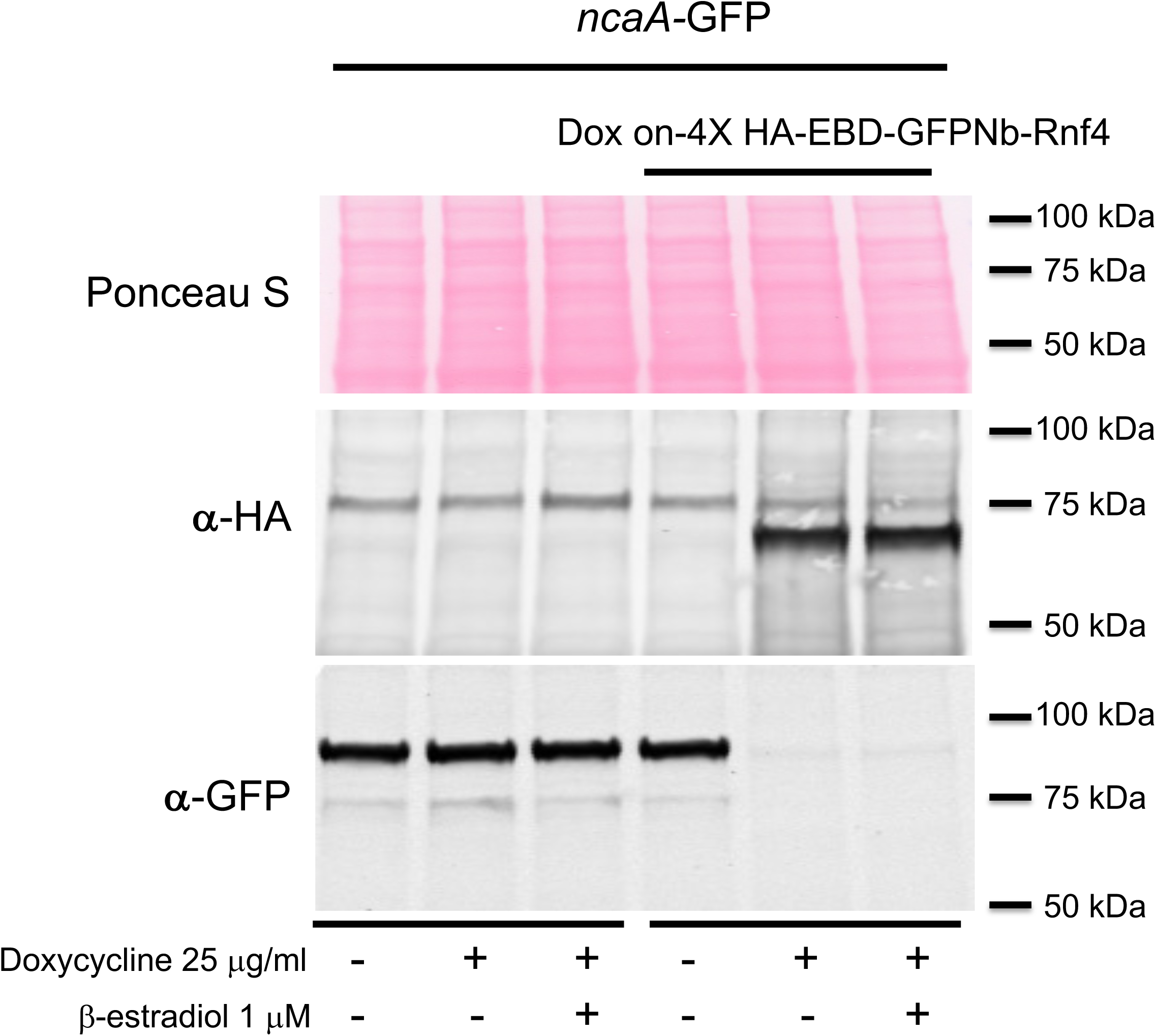
Degradation of a nuclear factor by GFPNb-directed proteolysis. A strain containing a *ncaA*-GFP fusion gene was transformed with the doxycycline-inducible GFPNb-Rnf4-producing fusion gene. The untransformed *ncaA*-GFP strain was used as a control. These strains were grown in Sabouraud dextrose media with or without doxycycline and β-estradiol as indicated at the bottom of the panel. Whole cell protein extracts were prepared and analyzed as described above. Molecular mass standards are shown on the right.

The full-length NcaA-GFP protein was readily visualized using anti-GFP antisera at 92 kDa. Induction of the EBD-GFPNb-Rnf4 fusion protein upon the addition of doxycycline led to elimination of immunoreactive NcaA-GFP by western blotting. Lack of detectable NcaA-GFP upon doxycycline addition prevented our ability to determine if β-estradiol had any effect on the function of EBD-GFPNb-Rnf4. Clearly, the nanobody-containing protein was able to trigger proteolysis of a nuclear substrate as well as the cytoplasmic GFP above.

### Investigating the substrate range of EBD-GFPNb-Rnf4 in *A. fumigatus* cells

We next constructed several additional GFP fusion genes to explore the range of substrate proteins that EBD-GFPNb-Rnf4 would be able to trigger the proteolysis when coexpressed in the same strain. The three different gene products we fused to GFP were the ER-localized Cyp51A enzyme (25), the plasma membrane-localized ATP-binding cassette transporter protein AbcG1 (aka Cdr1B or AbcC (26)) and the nuclear transcription factor AtrR (22, 23). These different fusion proteins were all expressed in a strain containing the doxycycline-inducible EBD-GFPNb-Rnf4 nanobody construct and tested for doxycycline triggered proteolysis as above.

Doxycycline treatment of cells expressing either AbcG1- or AtrR-GFP fusion proteins led to an approximately 90% reduction of immunoreactive GFP (Figure 4). The Cyp51A-GFP fusion showed only a modest reduction to 70% of the untreated sample. These data suggest that the ability of EBD-GFPNb-Rnf4 can be seen for proteins localized to multiple different subcellular regions, although there are differences in the extent of degradation seen.

**Figure 4.**
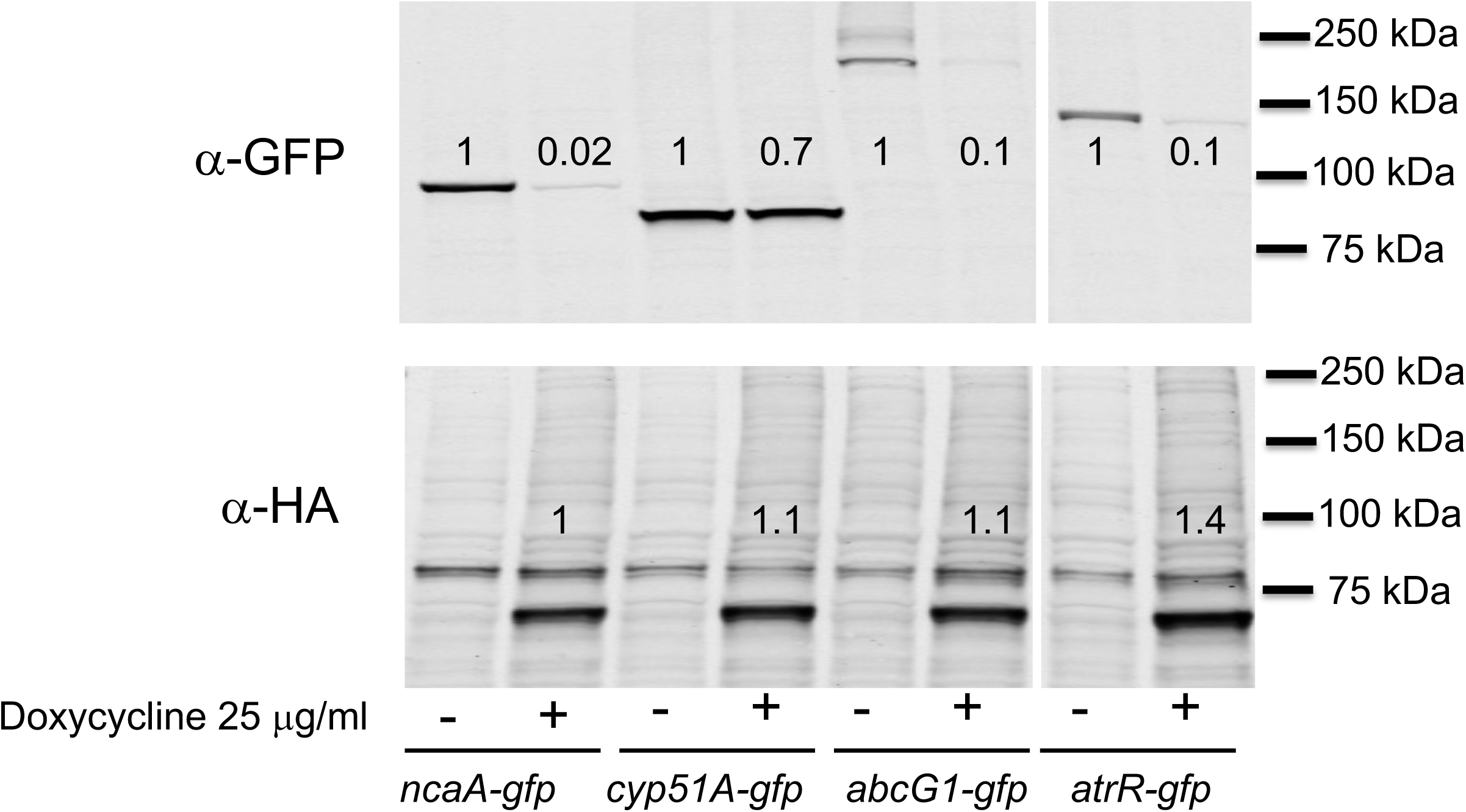
Comparative degradation of different GFP-tagged substrates. GFP cassettes were inserted at the carboxy-termini of 3 genes involved in azole susceptibility along with the previously characterized *ncaA* locus. These genes were *cyp51A*, *abcG1* and *atrR*. All these strains contained a copy of the doxycycline-inducible EBD-GFPNb-Rnf4-producing fusion gene integrated into their *fcyB* gene. Appropriate transformants were grown overnight in Sabouraud dextrose media with or without the addition of doxycycline and whole cell protein extracts prepared. Equal amounts of each extract were then analyzed by western blotting as above. Molecular mass standards are indicated and the numbers in each lane represent levels of protein normalized to the expression seen in the absence of doxycycline.

Since all of these proteins play a role in voriconazole susceptibility, we used a disc diffusion assay to test the impact of doxycycline on the resulting azole susceptibility phenotype. Conidia from the strains above were spread on minimal media containing (+) or lacking (-) doxycycline. A disc containing 1 μg of voriconazole was then placed in the center of the plate followed by incubation at 37°C. Plates were photographed after 48 hours (Figure 5).

**Figure 5.**
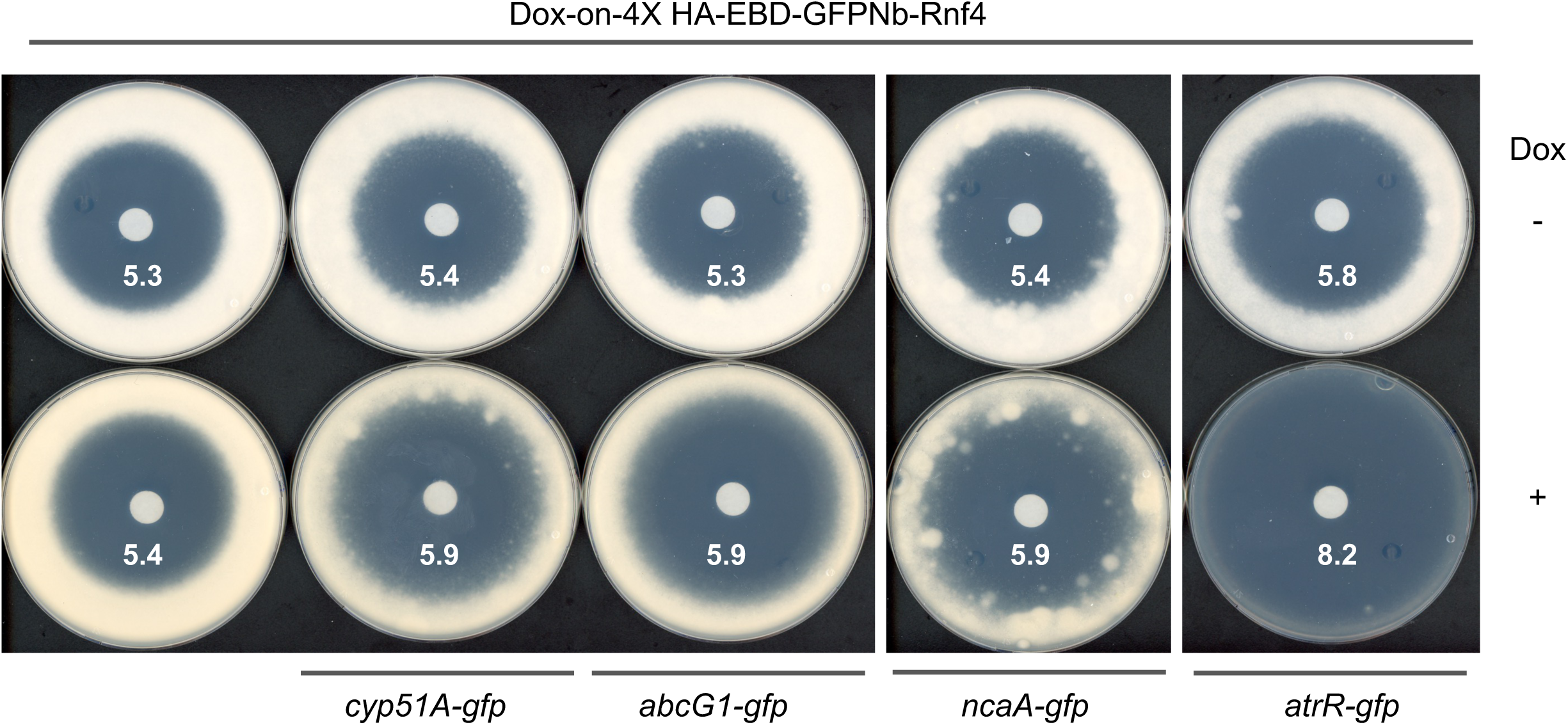
Doxycycline-inducible voriconazole phenotypes upon degradation of GFP fusion proteins. The doxycycline-inducible EBD-GFPNb-Rnf4-containing strain expressing GFP fusion genes corresponding to Cyp51A-, AbcG1-, NcaA- and AtrR-GFP were tested for their voriconazole phenotypes by a disc diffusion assay. ∼100 conidia were placed on Sabouraud dextrose plates containing (+) or lacking (-) 25 μg/ml doxycycline. A disk containing 1 μg of voriconazole was then placed in the center of each dish and incubated at 37°C for 48 hr. One representative assay of at least 3 trials is shown and the numbers refer to the diameter from the edge of the disk to the start of the growing fungal mat.

Doxycycline-induced proteolysis of AtrR-GFP caused the largest increase in voriconazole susceptibility while similar increases in voriconazole susceptibility were seen for Cyp51A-, AbcG1- and NcaA-GFP despite the differences in protein levels after doxycycline treatment. Possible reasons for these differences will be discussed below.

### Proteolysis of Erg6 and effects on gene regulation

Given that proteolysis of the ER-localized Cyp51A was only modestly efficient, we wanted to test a second protein localized to this compartment. We used Erg6 which has recently been shown to be an essential enzyme in *A. fumigatus* and localized to the ER (27). We also constructed a doxycycline-inducible GFPNb-Rnf4 fusion protein that lacked the EBD used above to directly evaluate any contribution of the EBD to proteolysis control. We constructed an *erg6*-GFP fusion in wild-type cells and nanobody-expressing cells. The three different *erg6*-GFP strains were grown in the presence (+) or absence (-) of doxycycline and then analyzed for levels of Erg6-GFP and the respective nanobody fusion protein by western blotting as before. A representative western blot is shown in Figure 6A.

**Figure 6.**
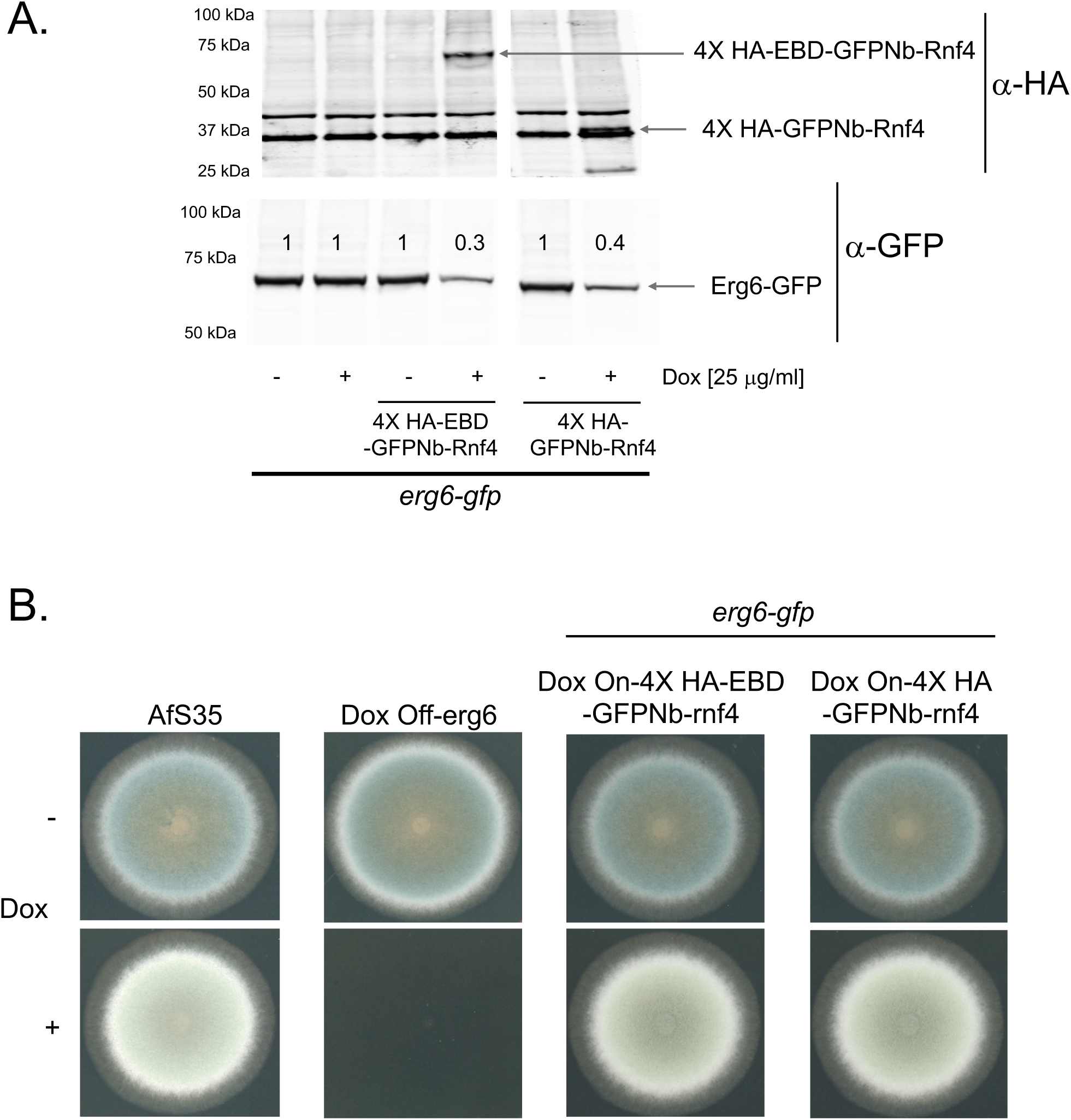
Regulated degradation of Erg6. A. Strains expressing Erg6-GFP fusion proteins along with doxycycline-inducible forms of either EBD-GFPNb-Rnf4 or GFPNb-Rnf4 were grown in the presence (+) or absence (-) of doxycycline and whole cell protein extracts prepared. Equal amounts of each extract were resolved on SDS-PAGE and analyzed by western blotting using the indicated anti-HA or anti-GFP antisera. Locations of the proteins of interest are denoted by the labeled arrows and molecular mass standards are listed at the left. B. Phenotypes caused by transcriptional repression of Dox off-erg6 or doxycycline-inducible proteolysis of Erg6-GFP were tested on media lacking (-) or containing (+) 25 mg/L doxycycline.

Doxycycline-dependent induction of either the EBD-GFPNb-Rnf4 or the GFPNb-Rnf4 nanobody fusion protein led to pronounced reduction in the level of Erg6-GFP. The molecular mass of each nanobody fusion protein was consistent with the predicted size of the GFPNb-Rnf4 with (EBD-GFPNb-Rnf4) or without (GFPNb-Rnf4) the EBD moiety.

Previous characterization of a doxycycline-repressible promoter fusion to *erg6* demonstrated that repression of *erg6* transcription caused both a growth defect but also an acute induction of ATP-binding cassette transporter-encoding gene expression (27). We obtained this fusion gene and introduced it into our wild-type strain background (AfS35) and constructed an isogenic doxycycline-repressible form of *erg6* (Dox off-erg6). As reported earlier using a different genetic background, Dox off-erg6 was not able to grow in the presence of doxycycline (Figure 6B) (27). Even though doxycycline induction of either nanobody construct in cells expressing Erg6-GFP caused proteolysis of this fusion protein (Figure 6A), this degree of protein loss did not prevent in vitro growth (Figure 6B). Given that Erg6-GFP levels were markedly reduced in nanobody-expressing cells, we wanted to test if a transcriptional response might be induced to permit growth maintenance in the face of reduced levels of this essential enzyme. We prepared total RNA from cells grown with or without doxycycline and used RT-qPCR to assess any transcriptional effect on selected genes.

We first examined levels of *erg6-gfp* mRNA and found that doxycycline-induced proteolysis induced a transcriptional increase of approximately two-fold in levels of *erg6-gfp* (Figure 7). We suspect that this increase in *erg6* transcription likely prevents a greater decrease in Erg6-GFP protein. We also assayed *abcG1*, *cyp51A* and *srbA* (encodes a key transcriptional regulator of ergosterol biosynthetic genes (28)) mRNA using appropriate primer pairs. Both *abcG1* and *cyp51A* were markedly elevated upon nanobody-dependent proteolysis of Erg6-GFP while a significant but lesser increase was observed for *srbA*. These data are consistent with the view that proteolysis of Erg6 caused altered ergosterol metabolism that leads to a transcriptional response.

**Figure 7.**
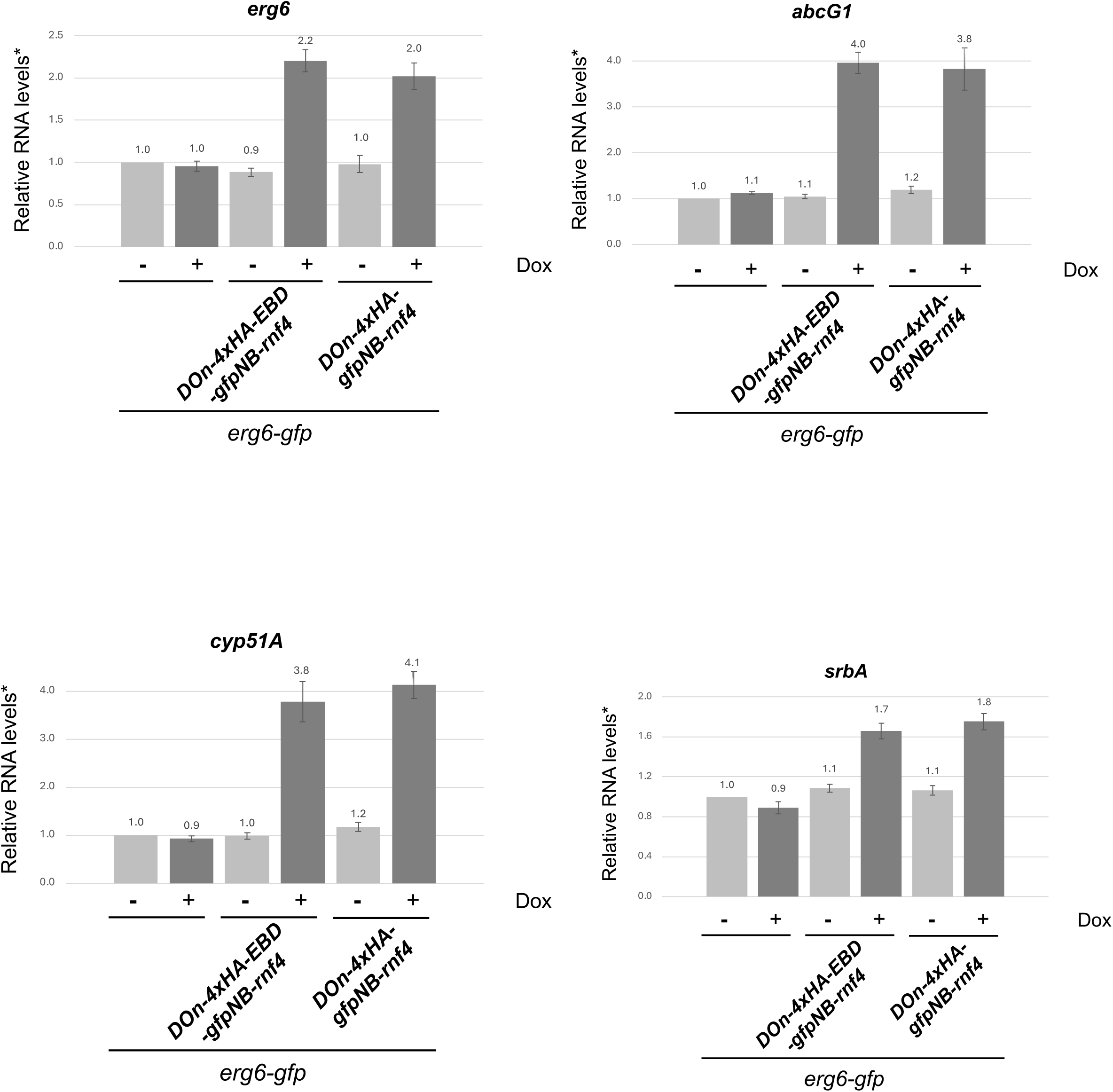
Transcriptional response to Erg6 proteolysis. Total RNA was prepared from the strains described above and analyzed by RT-qPCR using probes specific for *erg6*, *abcG1*, *srbA* or *cyp51A*. Levels of each transcript were normalized to those seen in wild-type cells with no doxycycline treatment.

## Discussion

### Nanobody-directed proteolysis is an effective means of protein depletion in Afu

While methods have already been described to vary production of mRNAs in Afu using regulated promoters, we demonstrate that expression of a GFP-directed nanobody coupled to an E3 ligase can effectively lower levels of GFP fusion proteins in this fungus. This provides an immediate reduction in levels of a desired target protein as opposed to the eventual depletion caused by a block in gene transcription. Our experiments to date have not been able to demonstrate any useful regulation with the inclusion of the EBD and future work will employ the small GFPNb-Rnf4 fusion protein that effectively causes regulated proteolysis.

The doxycycline-inducible promoter construct provides efficient control of expression of the GFPNb-Rnf4 fusion protein in which expression is strictly dependent on the presence of doxycycline. This is a crucial element for regulated degradation as production of the GFPNb-Rnf4 fusion protein determines the onset of proteolysis. While all the GFP fusion proteins exhibited some degree of proteolysis, the extent of this proteolysis varied. 3 of the 4 GFP fusion proteins were reduced to roughly 10% of their normal levels upon the addition of doxycycline but the Cyp51A-GFP fusion was still present at 70%, suggestive of this fusion protein being resistant to nanobody-mediated proteolysis. We do not think this is the case as while the level of Cyp51A-GFP remained relatively high, treatment of this strain with doxycycline caused a readily detectable increase in voriconazole susceptibility (Figure 5). The voriconazole zone of inhibition caused by doxycycline was very similar between Cyp51A-GFP, AbcG1-GFP and NcaA-GFP even though the reduction of the respective GFP fusion protein was quite different.

We interpret this to indicate that Cyp51A-GFP is impacted by expression of the nanobody and the resulting degradation triggers a compensatory response that masks the full effect caused by degradation of Cyp51A-GFP. Previous work comparing the voriconazole phenotype of a *cyp51Aϕλ* with *cyp51Bϕλ* demonstrated that loss of the Cyp51A protein caused a more pronounced increase in azole susceptibility than the corresponding loss of Cyp51B (25). We detected an increased voriconazole susceptibility upon degradation of Cyp51A to only 70% of its normal level while these previous experiments completely eliminated Cyp51A. These data support the view that Cyp51A has a unique importance in azole resistance in Afu, even in the presence of normal Cyp51B.

### Proteins can be targeted in different compartments

Given the heterologous nature of this inducible proteolysis system, we wanted to determine if Afu proteins with different subcellular distributions could be accessed by the GFPNb-Rnf4 fusion protein. Inducible production of GFPNb-Rnf4 led to the degradation of proteins localized to the cytoplasm (GFP), the endoplasmic reticulum (Cyp51A, Erg6), the nucleus (AtrR, NcaA) and the plasma membrane (AbcG1). Although not all targets were degraded to the same extent (as discussed above), all were susceptible to nanobody-mediated proteolysis.

This wide range of accessible locations for GFPNb-Rnf4 gives support for potential use of this system to produce a wide-range of proteolytically-susceptible substrates such as might be generated in a genome-wide collection of GFP fusions. Similar strategies have been used in *Saccharomyces cerevisiae* (29–31). These parallel approaches typically include use of the auxin-inducible degron which we found to be problematic to deploy in Afu. The previous *S. cerevisiae* technologies and the one we describe here have the advantage of being able to acutely deplete even essential proteins to allow their functional analysis over appropriate time courses. Construction of a GFP fusion library of Afu is an important future goal and use of the GFPNb-Rnf4-expressing strain would allow this library to be used for systematic depletion of any desired GFP fusion protein.

### Nanobody-mediated depletion can generate hypomorphic alleles

While chronic disruption alleles have been valuable genetic reagents permitting study of the roles of Afu genes, we found that nanobody-dependent proteolysis of GFP fusion proteins was able to produce a wider range of function than we initially expected. In simple cases (AbcG1, AtrR, NcaA), their respective GFP fusions behave as loss-of-function alleles upon induction of nanobody expression with the level of each protein reduced to roughly 10% or less of wild-type. In the case of Erg6-GFP, we can deplete this protein with GFPNb-Rnf4 expression but, even in the presence of doxycycline and this depletion of Erg6 activity, the cells still grow. This is very different from the behavior seen when a doxycycline-repressible promoter is used to drive transcription of the essential *erg6* gene (27). The presence of doxycycline prevents growth of a doxycycline-repressible *erg6* gene. This indicates that, at least for *erg6*, the GFPNb-Rnf4-mediated degradation of Erg-GFP does not mimic a loss-of-function allele.

Analysis of the mRNA levels of several different genes linked to activity of the ergosterol biosynthetic pathway indicated that proteolysis of Erg6-GFP gave rise to a signal that engaged a compensatory response to loss of this essential enzyme. Proteolysis of Erg6-GFP led to an induction of *erg6*-GFP and *cyp51A* mRNA, with a coordinate rise in *srbA* transcript. The reduction in Erg6 activity was sufficient to trigger elevation of the expression of another ergosterol pathway enzyme as well as the transcription factor-encoding gene *srbA* (28). Note that the observed elevation of *erg6*-GFP is most likely to due to the transcriptional activation of the native erg6 promoter in this construct. Previous work using a doxycycline-repressible promoter in place of wild-type *erg6* transcriptional signals could not detect a compensatory feedback response upon reduction of *erg6* mRNA (27). Additionally, the elevation of *cyp51A* suggests multiple genes within the ergosterol biosynthetic pathway are activated upon Erg6-GFP proteolysis, possibly via increased SrbA activity.

These data support our view that the *erg6*-GFP is a conditional hypomorphic allele depending on expression of the GFPNb-Rnf4. While the previously constructed doxycycline-repressible *erg6* allele behaves as a null version of *erg6* in the presence of doxycycline, the acute degradation of Erg6-GFP triggers a transcriptional response of the ergosterol pathway to adjust for the lowered levels of the Erg6 enzyme. This behavior illustrates a valuable property of the nanobody-mediated proteolysis of a targeted gene product. Some fusion proteins (AbcG1, AtrR) are readily degraded as GFP fusions and we did not detect any transcriptional response to correct their loss.

Degradation of the Erg6-GFP, with its likely reduction in ergosterol biosynthetic pathway function, was sufficient to induce a transcriptional correction. This nanobody system provides both a means to directly deplete a protein of interest but also to generate allelic variants that can uncover transcriptional circuitry. We anticipate this approach will be useful for future analyses of genes in *A. fumigatus*.

## Material and Methods

### *fumigatus* strains, media and transformation

All strains used in this study were derived from the AfS35 (FGSC #A1159) strain. *A. fumigatus* strains were typically grown at 37°C in rich medium (Sabouraud dextrose; 0.5% tryptone, 0.5% peptone, 2% dextrose [pH 5.6 ± 0.2tt). Selection of transformants were made in minimal medium (MM; 1% glucose, nitrate salts, trace elements, 2% agar [pH 6.5tt); trace elements, vitamins, and nitrate salts are as described in the appendix of reference (32), supplemented with 1% sorbitol. For selection of GFPNb-*rnf4* strains, that are targeted to the *fcyB* transporter-encoding gene as described (33), unbuffered minimal medium (∼pH 4.5) was supplemented with 2.5 mg/L 5-flucytosine (5-FC). For selection of phleomycin-resistant, GFP-expressing transformants, minimal media was buffered to pH 7 with 10M NaOH and supplemented with 20 mg/L phleomycin. For solid medium, 1.5% agar was added. Dox off promoter shut off experiments were performed by adding 25 mg/liter doxycycline (BD Biosciences).

All strains used in this study are listed in table 1. GFPNb-*rnf4* strains were generated as listed below. Initially, the 4X HA-EBD-GFPNb-*rnf4* (construct 1) was synthesized by Genscript Inc. and cloned into DraIII-SacII digested pϕλfcyB-tetON-sGFP (kindly provided by Fabio Gsaller (34)) to make the plasmid pSP141. Sequences included in construct 1 were: EBD (human estrogen binding domain: LEPSAGDMRAANLWPSPLMIKRSKKNSLALSLTADQMVSALLDAEPPILYSEYDPTRPFSEASMMGLLTNLADRELVHMINWAKRVPGFVDLTLHDQVHLLECAWLEILMIGLVWRSMEHPGKLLFAPNLLLDRNQGKCVEGMVEIFDMLLATSSRFRMMNLQGEEFVCLKSIILLNSGVYTFLSSTLKSLEEKDHIHRVLDKITDTLIHLMAKAGLTLQQQHQRLAQLLLILSHIRHMSNKGMEHLYSMKCKNVVPLYDLLLEMLDAHRLHAPTSRGGASVEETDQSHLATAG STSSHSLQKYYITGEAEGFPATV), GFP nanobody (single chain anti-GFP-antibody derived from a camel single-heavy chain antibody; sequence: VQLVESGGALVQPGGSLRLSCAASGFPVNRYSMRWYRQAPGKEREWVAGMSSAGDRSSYEDSVKGRFTISRDDARNTVYLQMNSLKPEDTAVYYCNVNVGFEYWGQGTQVTV SS) and Rnf4 (E3 ubiquitin ligase ring finger protein 4 from rat; sequence: ASEERRRPRRNGRRLRQDHADSCVVSSDDEELSKDKDVYVTTHTPRSTKDEGTTGLRPSGTVSCPICMDGYSEIVQNGRLIVSTECGHVFCSQCLRDSLKNANTCPTCRKKINHKRYHPIYIGSGTVSCPICMDGYSEIVQNGRLIVSTECGHVFCSQCLRDSLKNANTCPTCRKKINHKRYHPIYI) that were all codon optimized for expression in *A. fumigatus* to generate construct 1. The EBD was flanked by AfeI sites to allow eviction of this domain. Plasmid pSP141 was digested with HindIII and SacII to release the 4.2kb tetTA-tetO (hereafter referred to as Dox on or DO)-4X HA-EBD-GFPNb-rnf4 (construct 2) and cloned into fcyB-MH-CO F.luc_pUC57kan to replace firefly luciferase with construct 2 to generate plasmid pSP148. This plasmid has construct 2 flanked by 50 bp homology to the upstream and downstream regions of the *fcyB* gene. Plasmid pSP150 was generated by removing the EBD by AfeI digestion and subsequent self-ligation of the plasmid by T4 DNA ligase. The plasmid sequence of pSP148 and pSP150 can be accessed (To be deposited in Addgene). Transforming fragments from pSP148 and pSP150 were generated by digestion with PacI and PmeI to allow targeting to *fcyB* and production of strains SPF367 (DO-4X HA-EBD-GFPNb-*rnf4*) and SPF393 (DO-4X HA-GFPNb-*rnf4*), respectively. Homologous targeting was accomplished using microhomology-mediated CRISPR-Cas9 directed insertion at the *fcyB* locus using fcyB upstream (5’-ctcaacgaccaggtaaggtgagg) and fcyB downstream (5’-ctcttgatgatagaagtgtgcgg) crRNAs. This led to the deletion of native *fcyB* and acquisiton of 5-FC resistance. Strains expressing GFP fusion proteins were generated in either SPF367 or SPF393 using PCR cassettes containing a GA5 linker-GFP-Ble (phleomycin resistance marker) from gfp2.5-pSK494-pRFP (procured from the Fungal Genetics Stock Centre) using primers containing 50 bp at the end of the respective coding sequence and 50 bp at the 3’ end of the gene to be studied. The Afu codon optimized GFP sequence used was: atgtccaagggcgaggaactgttcaccggcgtcgtccctatcctggtcgagctggacggtgacgtcaacggtcacaagttctccgtcagcggcgagggcgagggcgacgccacctacggcaagctgaccctgaagttcatctgcaccaccggcaagctgcccgtcccttggcccaccctggtcaccaccttcacctacggcgtccagtgcttctcccgctaccccgaccatatgaagcgccacgacttcttcaagtccgccatgcccgagggctacgtccaagagcggaccatcttcttcaaggacgacggcaactacaagacccgcgctgaggtcaagttcgagggtgacaccctggtcaaccgcatcgagctgaagggcatcgacttcaaggaagatggcaacatcctgggccacaagctcgagtacaactacaactcccacaacgtctacatcatggccgacaagcagaagaacggcatcaaggccaacttcaagactcggcacaacatcgaggacggcggcgtccagctggccgaccactaccagcagaacacccccatcggcgacggccccgtcctgctgcccgacaaccactacctgtccacccagtccgccctgtccaaggaccccaacgagaagcgcgaccacatggtcctgctcgagttcgtcaccgccgctggcatcacccacggcatggacgagctgtacaagtga. Phleomycin resistant transformants were confirmed for targeted integration by validating the presence of novel junctions formed by GFP tagging of the gene by PCR as well as by checking the presence of the GFP-tagged protein by Western blotting with anti-GFP antibody.

### Drug disc diffusion/radial growth assay

Fresh spores of *A. fumigatus* were suspended in 1x phosphate-buffered saline (PBS) supplemented with 0.01% Tween 20 (1x PBST). The spore suspension was counted using a hemocytometer to determine the spore concentration. Spores were then appropriately diluted in 1x PBST. For the drug diffusion assay, 1×10^6^ spores were mixed with 10 ml soft agar (0.7%) and poured over 15 ml regular agar (1.5%) containing Sabaraud dextrose medium. A paper disk was placed on the center of the plate, and 10 ml of either 1 mg/liter voriconazole was spotted onto the sterile filter paper. For the radial growth assay, ∼100 spores (in 4 μl) were spotted on Sabaraud dextrose medium in the presence or absence of 25 mg/ml doxycycline. The plates were incubated at 37°C and scanned after 2 days.

**TABLE 1.**
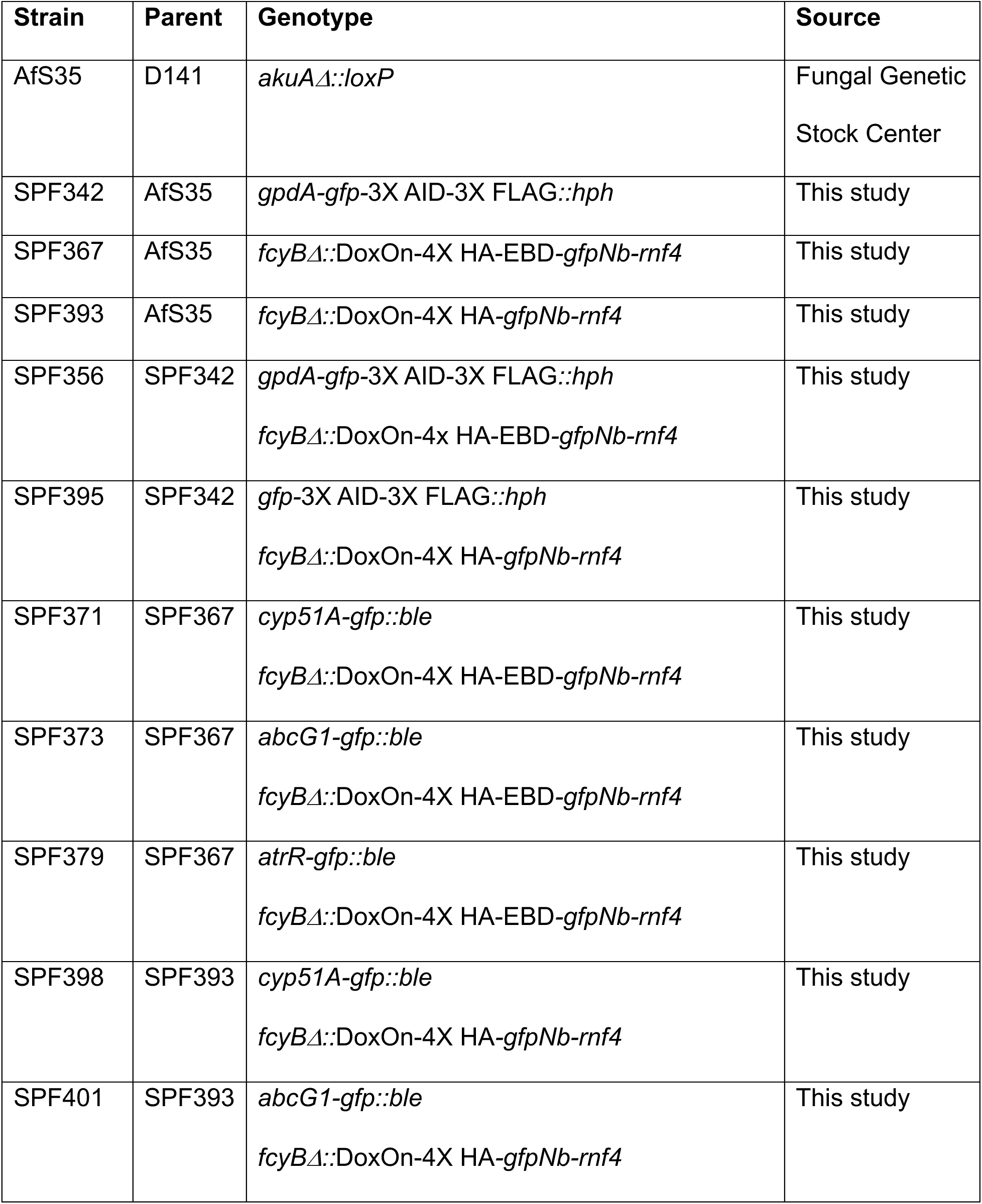
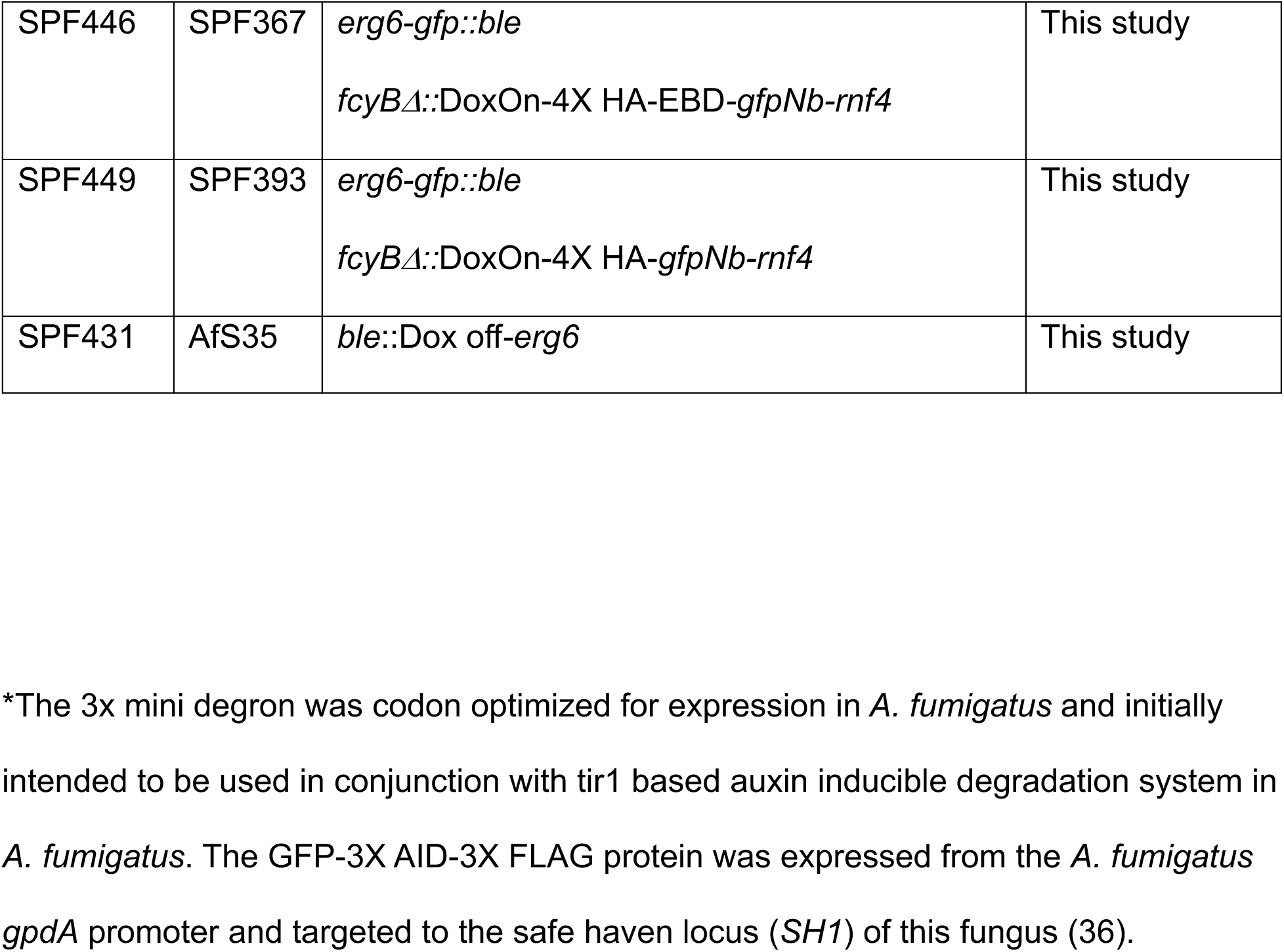
*A. fumigatus* strains used in this study.

| Strain | Parent | Genotype | Source |
| --- | --- | --- | --- |
| AfS35 | D141 | *akuAΔ::loxP* | Fungal Genetic<br>Stock Center |
| SPF342 | AfS35 | *gpdA-gfp-3X AID-3X FLAG::hph* | This study |
| SPF367 | AfS35 | *fcyBΔ::DoxOn-4X HA-EBD-gfpNb-rnf4* | This study |
| SPF393 | AfS35 | *fcyBΔ::DoxOn-4X HA-gfpNb-rnf4* | This study |
| SPF356 | SPF342 | *gpdA-gfp-3X AID-3X FLAG::hph*<br>*fcyBΔ::DoxOn-4x HA-EBD-gfpNb-rnf4* | This study |
| SPF395 | SPF342 | *gfp-3X AID-3X FLAG::hph*<br>*fcyBΔ::DoxOn-4X HA-gfpNb-rnf4* | This study |
| SPF371 | SPF367 | *cyp51A-gfp::ble*<br>*fcyBΔ::DoxOn-4X HA-EBD-gfpNb-rnf4* | This study |
| SPF373 | SPF367 | *abcG1-gfp::ble*<br>*fcyBΔ::DoxOn-4X HA-EBD-gfpNb-rnf4* | This study |
| SPF379 | SPF367 | *atrR-gfp::ble*<br>*fcyBΔ::DoxOn-4X HA-EBD-gfpNb-rnf4* | This study |
| SPF398 | SPF393 | *cyp51A-gfp::ble*<br>*fcyBΔ::DoxOn-4X HA-gfpNb-rnf4* | This study |
| SPF401 | SPF393 | *abcG1-gfp::ble*<br>*fcyBΔ::DoxOn-4X HA-gfpNb-rnf4* | This study |
| SPF446 | SPF367 | *erg6-gfp::ble*<br>*fcyBΔ::DoxOn-4X HA-EBD-gfpNb-rnf4* | This study |
| SPF449 | SPF393 | *erg6-gfp::ble*<br>*fcyBΔ::DoxOn-4X HA-gfpNb-rnf4* | This study |
| SPF431 | AfS35 | *ble::Dox off-erg6* | This study |

\*The 3x mini degtron was codon optimized for expression in *A. fumigatus* and initially intended to be used in conjunction with tir1 based auxin inducible degradation system in *A. fumigatus*. The GFP-3X AID-3X FLAG protein was expressed from the *A. fumigatus* *gpdA* promoter and targeted to the safe haven locus (*SH1*) of this fungus (36).

### Western blotting

Western blotting was performed as described in (35) with the following modifications. The GFP (B-2) monoclonal antibody (sc-9998; Santa Cruz Biotechnology) was used at a 1:1000 dilution while the HA monoclonal antibody (2-2.2-14; Invitrogen) was used at 1:2500 dilution.

### Measurement of mRNA level

Reverse transcription quantitative PCR (RT-qPCR) was performed as described in reference (24), with the following modification. Cell lysates were prepared from mycelial biofilm cultures formed upon inoculating 10^6^ spores in a petri dish containing 20 mL of Sabouraud dextrose broth, either in the presence or absence of doxycycline (25 mg/L) and allowing growth for 24 h at 37°C under nonshaking conditions. The Ct value of the gene coding for *tef1* was used for normalization of variable cDNA levels to determine the fold difference in transcript levels.

## Acknowledgements

We thank Drs. Jarrod Fortwendel, Damian Krysan, Rob Piper and Scott Filler for helpful discussions. Important materials were provided by Drs. Jarrod Fortwendel and Fabio Gsaller.

